# Simple, low-cost, and well-performing method, the outgrowth technique, for the isolation of epithelial cells from nasal polyps

**DOI:** 10.1101/2023.01.10.522992

**Authors:** Jonghui Kim, Karla Hegener, Claudia Hagedorn, Kaschin Jamal Jameel, Daniel Weidinger, Inga Marte Charlott Seuthe, Sabine Eichhorn, Florian Kreppel, Jonas Jae-Hyun Park, Jürgen Knobloch

## Abstract

**Objectives:** Epithelial cells are an important part of the pathomechanism in chronic rhinosinusitis with nasal polyps. It is therefore essential to establish a robust method for the isolation and culture of epithelial cells from nasal polyps to enable further research. In this study, the feasibility of the outgrowth technique for the isolation of the epithelial cells from the nasal polyps was evaluated.

**Methods:** The outgrowth technique was performed to isolate the epithelial cells. Proliferation was evaluated up to the 3rd passage. Epithelial cells were identified and differentiation and proliferation were evaluated using flow cytometry with anti-cytokeratin, anti-p63, and anti-Ki-67. A functionality test was assessed by determining type 2–relevant proteins using ELISA, representatively, interleukin-33 and periostin.

**Results:** Using the outgrowth technique, epithelial cells could be isolated from all tissue samples. Isolated epithelial cells showed a proliferation rate of approximately 7- to 23-fold every 6 days up to the 3rd passage. Over 97% of isolated cells were shown to be cytokeratin- and p63-positive, and over 86% of them were Ki-67–positive in flow cytometry. Interleukin-33 and periostin were detectable in the supernatant.

**Conclusions:** We introduce a simple, low-cost, and well-performing method for isolating epithelial cells from nasal polyps with the outgrowth technique.

## INTRODUCTION

Nasal polyps can be developed in the course of chronic rhinosinusitis, which is characterized as the inflammation of the mucous of the nasal and paranasal sinuses. Chronic rhinosinusitis with nasal polyps (CRSwNP) is generally characterized as type 2 inflammation in western countries [1]. Much evidence, such as the infiltration of eosinophilic granulocytes in nasal polyps, has suggested that CRSwNP is a type 2 inflammatory disease [1]. Furthermore, the recently approved therapies with dupilumab (anti-interleukin-4/anti-interleukin-13), mepolizumab (anti-interleukin-5), and omalizumab (anti-immunoglobulin E) in patients with nasal polyposis are highly effective [2–4], so it is strongly indicated that the regulation of interleukin-4 (IL-4), interleukin-5 (IL-5), and immunoglobulin E (IgE) via the type 2– inflammation track may play a crucial role in the pathogenesis.

The epithelial cells seem to be an important part of the entire pathomechanism of CRSwNP [5]. The epithelial cells of nasal polyps are a distinct pathological cell type and differ from healthy nasal epithelial cells. As such, basal cell hyperplasia, features of epithelial-mesenchymal transition, and deregulation of epithelial differentiation markers and of key regulators of neurogenic inflammation are indicators for pathological epithelial cells in nasal polyps [6–8]. Epithelial cells also play an important role in type 2 inflammation by secreting a large number of pro-inflammatory mediators, such as interleukin-33 (IL-33) [9], which on the one hand activate type 2 innate lymphoid cells (ILC2) and mast cells directly and, on the other hand, support the conventional dendritic cell–mediated activation of Th2 cells and the subsequent production of IgE produced by B cells [5]. As a result, remodeling takes place, with a strong change in the composition of the epithelium, resulting, for example, in basal cell hyperplasia, ciliary dysfunction, and impaired mucus production of the epithelial cells [5]. The mechanism of the remodeling process in CRSwNP is, however, not fully understood, but it might be an auspicious drug target. Therefore, it is mandatory to establish a preclinical method, particularly a cell culture model enabling a wide range of investigations to better understand the role of epithelial cells in CRSwNP.

Here, we introduce a simple, low-cost, and well-performing method for isolating and cultivating epithelial cells from nasal polyps by using an outgrowth technique without enzymatic pretreatment.

## MATERIALS AND METHODS

### Ethics statement and patient materials

This study was performed according to the ethical standards of the declaration of Helsinki and following approval of the ethical committees of the university of Witten/Herdecke, Germany (no. 209/2020). Donors gave informed consent for the research use of their nasal polyp tissue. Pseudonymisation was carried out, and patient data were treated confidentially.

Nasal polyps obtained from patients with CRSwNP and undergoing endoscopic endonasal sinus surgery are the source of the tissue material. We performed a preoperative clinical endoscopic examination and a computed tomography scan of sinus. The patients reported typical symptoms of chronic rhinosinusitis, such as rhinorrhea, cephalgia, and obstructed nasal breathing. Patients with a unilateral finding were excluded due to possible neoplasia and papilloma.

We include tissues from 6 patients for this study (Table 1). The evaluation of the isolation, proliferation, and flow cytometric analysis was carried out with Polyp 1, Polyp 2, and Polyp 3. ELISA was performed with epithelial cells from Polyp 4, Polyp 5, and Polyp 6.

**Table 1.** The demographic and clinical characteristics of tissue doners in this study.

|  | **Polyp 1** | **Polyp 2** | **Polyp 3** | **Polyp 4** | **Polyp 5** | **Polyp 6** |
| --- | --- | --- | --- | --- | --- | --- |
| **Age** | 45 | 65 | 45 | 47 | 62 | 58 |
| **Sex** | Female | Male | Male | Male | Female | Male |
| **Nasal polyp score**<br>**(right/left)** | 2/3 | 3/1 | 3/4 | 2/2 | 2/2 | 3/3 |
| **Prior sinus surgeries** | No | No | No | No | No | Yes |
| **Tissue eosinophilia** | Yes | Yes | Yes | Yes | Yes | Yes |
| **Asthma** | No | No | No | No | Yes | No |
| **NSAID-exacerbated**<br>**respiratory disease** | No | No | Yes | No | Yes | No |

### Isolation of epithelial cells from nasal polyp tissue

All isolation and culture procedures were performed in a sterile condition with sterile instruments. Tissues were rinsed with 7.5% povidone-iodine solution (Braunol^®^, B.Braun, Germany) and rinsed twice with 0.9% sodium chloride immediately after harvest from sinus surgery. The tissue was stored in Dulbecco’s Modified Eagle Medium (DMEM; Gibco, cat #: 1437676, Carlsbad, CA, USA) with 10% fetal bovine serum (v/v; Bio&SELL, cat #: FBS.S 0615 HI, Feucht, Germany) and 1% penicillin/streptomycin (Bio&SELL, cat #: BS.A 2213, Feucht, Germany) at 4°C. The following treatment was carried out within 6 hours after surgical removal. The tissue was washed once with Dulbecco’s phosphate-buffered saline without calcium and magnesium (PBS; Gibco, cat #: 14190169, Carlsbad, CA, USA). Polyps were cut into small pieces on the petri dish with surgical scissors (approx. 1–1.5 mm diameter) (Figure 1A). The small pieces were placed in the 12-well culture plate (2 pieces/well) and placed under the safety cabinet without lid for 5–10 min to dry and stick to the bottom of the culture dish. Subsequently, 350 μl of Bronchial Epithelial Growth Medium (BEGM; Lonza, cat #: CC-3170, Basel, Switzerland) was added carefully without mobilizing the tissue pieces from the dish (Figure 1B); the wells should be completely covered with medium, but the medium volume must be as low as possible to avoid floating of the tissue pieces in the cell culture vessel (approx. 350 μl medium in our conditions, as mentioned above). The medium was renewed every 2 days without mobilizing the tissue pieces. The incubation lasted for 8 days, and the outgrowth of the epithelial cells from polyp tissues was evaluated microscopically every 2 days when the medium was changed. The cells were treated with 300 μl trypsin 0.25%/EDTA 0.02% for 2–3 min after rinsing with PBS twice, and the trypsin was neutralized with the same volume of soybean trypsin inhibitor (1 mg/ml in PBS; Sigma, cat #: SLCF8902, Steinheim, Germany). The cells were subsequently washed and centrifuged twice with PBS (300 × g) and then seeded on a 6-well plate (5,000 cells per well with 1.5 ml BEGM). The cells were grown in a humidified incubator with 5% CO_2_ at 37°C. The medium was renewed every 2 days. Cells were trypsinized at 70%–80% confluence and were re-passaged. The cell count was determined with the haemocytometer after mixing cell suspension with 0.1% trypan blue 1:1 (v/v) to identify vital cells.

**Figure 1.**
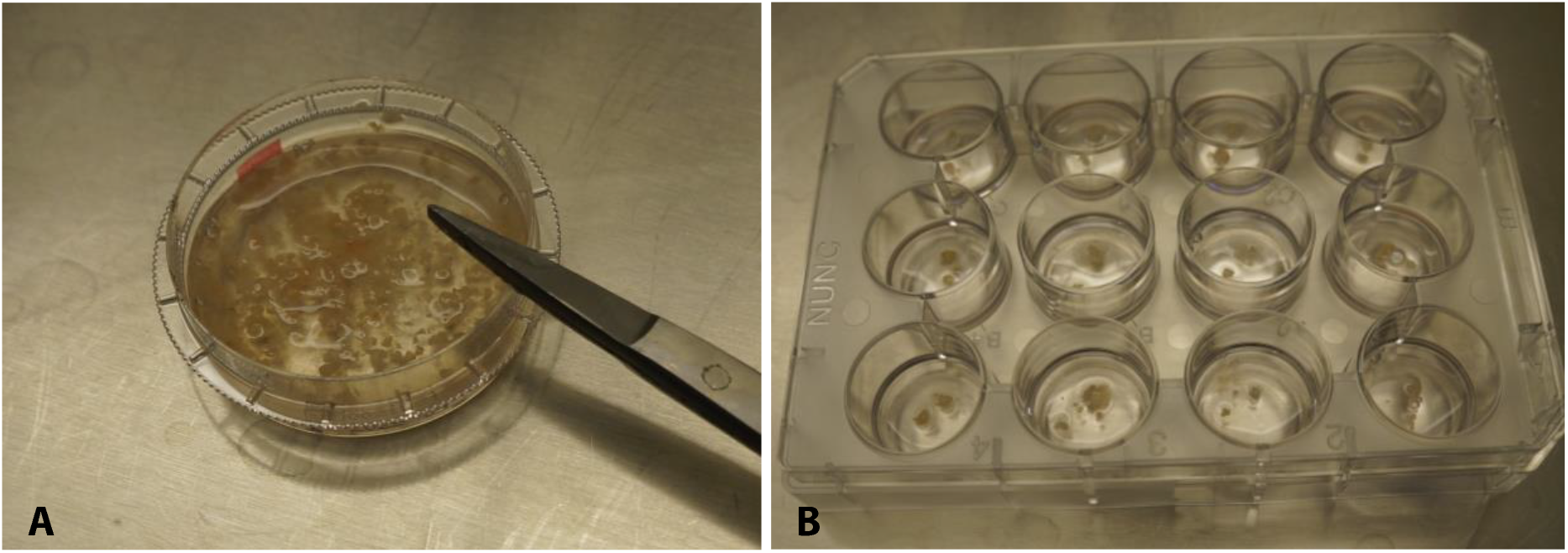
Tissue is on the petri dish placed and 1 ml PBS is added to keep the tissue from becoming dry (A). The tissue is cut into small pieces of approx. 1-1.5 mm with surgical scissors. Small pieces were placed on the 12-well culture plate (2 pieces/well) and stand under the safety cabinet without lid for 5-10 minutes to dry and stick on the surface of the culture dish. Approx. 350 μl of BEGM was then given carefully without mobilizing tissue pieces from the dish (B).

### Flow cytometry

Flow cytometry with anti-cytokeratin-PE (Miltenyi Biotec, clone: REA831, Bergisch Gladbach, Germany) and anti-p63 (Ventana Medical, clone: 4A4, AZ, USA) was performed to identify epithelial cells and evaluate differentiation. Simultaneously, the expression of Ki-67 (anti-Ki-67-FITC, Miltenyi Biotec, clone: REA183, Bergisch Gladbach, Germany) was determined to evaluate proliferation. Cells from the 2nd or 3rd passage were used for the analysis. Cells were cultured in a 75 cm^2^ cell culture flask with 10 ml BEGM, renewing the medium every 2 days. In 300 μl incubation buffer (0.5% BSA in PBS), 5 × 10^5^ - 1 × 10^6^ cells were collected. Supernatant was removed after centrifuging at 300 × g for 10 min at 4°C, and the cells were washed 3 times in total in the same way, by adding 300 μl incubation buffer followed by centrifugation. (The same washing process was carried out after each step and will not be mentioned from here on.) Cells were fixed in 500 μl of 4% formaldehyde for 15 min at room temperature. After removing the formaldehyde, permeabilization was carried out with 300 μl precooled permeabilization buffer (0.5% Octoxynol-9, 0.68 μM ethylenediaminetetraacetic acid, and 1% bovine serum albumin in distilled aqua) for 10 min on ice. The permeabilization buffer was removed, and cells were incubated first with 300 μl unconjugated anti-p63 (dilution 1:200 in incubation buffer) overnight at 4°C and then with secondary anti-mouse AlexaFluor633-conjugated antibody (dilution 1:200 in incubation buffer; Molecular Probes, cat #: A-21052, Eugene, OR, USA) for 1 h in a darkened chamber at room temperature. Cells were washed and incubated with fluorescent-dye conjugated anti-cytokeratin-PE and anti-Ki-67-FITC (each dilution 1:200 in incubation buffer) overnight at 4°C in a darkened chamber. Cells were washed and resuspended in 100 μl incubation buffer to perform a flow cytometric analysis. Compensation was set up before analysis, and positive and negative populations were predefined accordingly.

### Enzyme-linked immunosorbent assay (ELISA)

Epithelial cell function was assessed by the production of typical cytokines of type 2 inflammation, representatively IL-33 and periostin. Concentrations of IL33 and periostin in culture supernatants were measured by ELISA according to the manufacturer’s instructions (Invitrogen, cat #: BMS2048TEN, Carlsbad, CA, USA; R&D Systems, cat #: DY3548B, Minneapolis, MN, USA) and as described previously [10]. To obtain the supernatants, 5,000 cells were seeded in a 24-well plate with 1 ml BEGM, and medium was renewed every 2 days until 70%–80% confluence. Then 1 ml of BEGM was added, and supernatants were collected after 24, 48, and 72 hours. Cells from 2nd or 3rd passage were used for the analysis.

## RESULTS

### Isolation of nasal epithelial cells from polyp tissue, using the outgrowth technique, and evaluation of proliferation

Epithelial cells from the polyp tissue were observed by culturing using the outgrowth technique. First, outgrowth from the tissue samples of 3 patients was evaluated. Per sample, 24 small pieces of tissue were taken, as described above, to isolate epithelial cells. The outgrowth of epithelial cells was evaluated every 2 days. Almost all small pieces of tissue shed epithelial cells within 8 days (Figure 2): 19 of 24 tissue pieces from Polyp 1 yielded cells, and epithelial cells grew out from all 24 tissue pieces of Polyp 2 and Polyp 3. It was possible to isolate the epithelial cells from all 3 samples (Figure 3).

**Figure 2.**
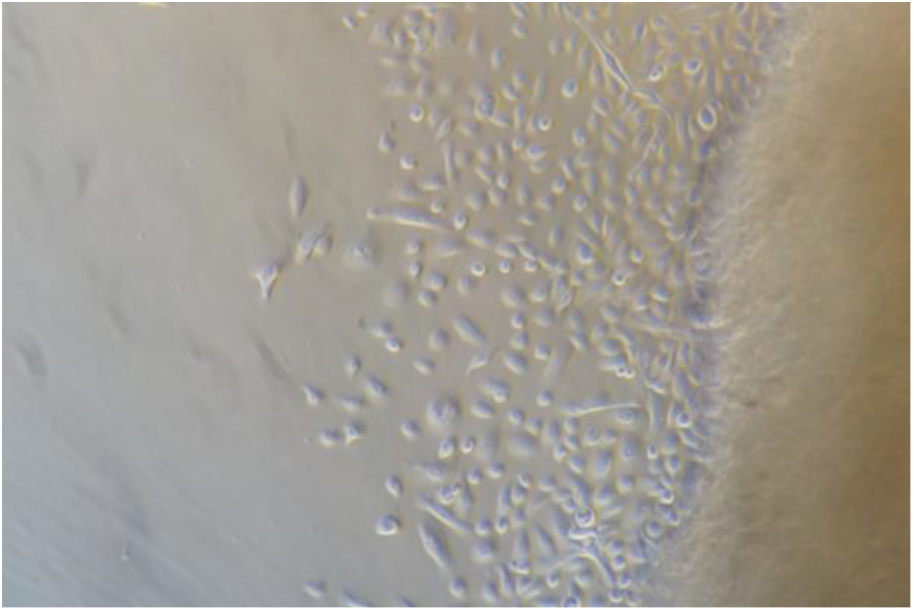
Epithelial cells grew out of polyp tissue. Outgrowth was observed in most tissues (19/24) in the first 8 days.

**Figure 3.**
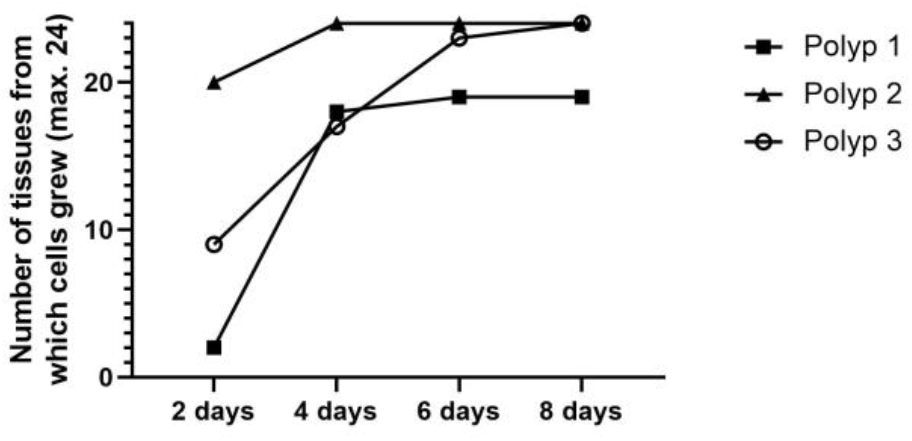
Cumulative number of tissue pieces from which cells grew out. Tissues from three patients were taken for isolation of epithelial cells. 24 small-cut pieces of tissue per sample from each patient (two pieces per well in a 12-well plate) were used for outgrowth of epithelial cells. The cumulative number of tissue samples from which cells grew out was assessed every 2 days. Cells grew out from almost all tissue pieces (67/72) within 8 days – 19 of 24 tissue pieces from Polyp 1 yielded cells. Epithelial cells grew out from all tissue pieces of Polyp 2 and Polyp 3.

On Day 8 of outgrowth, 320,000 (Polyp 1), 90,000 (Polyp 2), and 320,000 (Polyp 3) cells were collected, respectively. Cells were transferred to a 6-well plate at 5,000 cells per well, and the number of cells from 6 wells together was counted every 6 days up to the 3rd passage. Epithelial cells showed an excellent proliferation rate of approximately 7- to 23-fold up to the 3rd passage (Figure 4).

**Figure 4.**
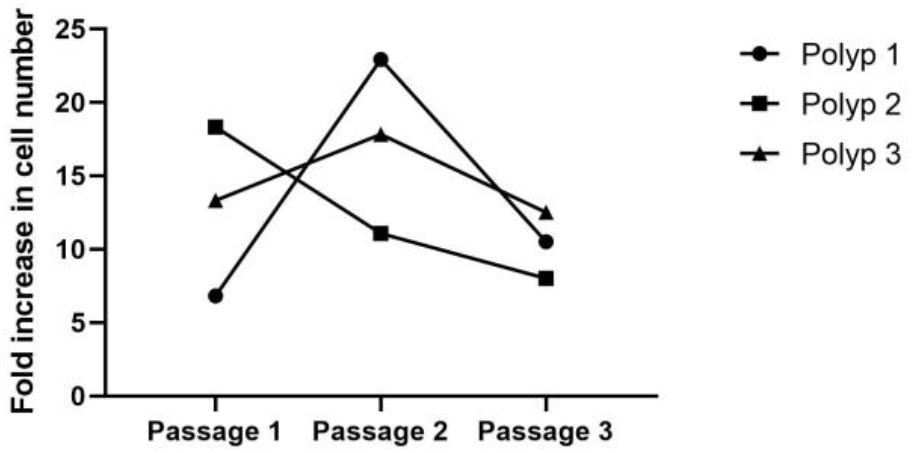
Fold increase in cell number up to the 3rd passage. Per well, 5,000 cells were seeded in a 6-well plate with 2 ml BEGM (30,000 cells in total). The medium was renewed every 2 days. The number of total cells from the 6-well plate was counted after 6 days. Epithelial cells showed a good proliferation rate, up to the 3rd passage, of approximately 7- to 23-fold. On Day 6 of each passage, approximately 205,000, 687,500, and 315,000 cells were counted from the Polyp 1 tissue; approximately 550,000, 332,500, and 240,000 cells from Polyp 2 tissue; and approximately 400,000, 535,000, and 375,000 cells from Polyp 3 tissue, respectively.

### Flow cytometric analysis to identify epithelial cells and evaluate differentiation and proliferation

To identify epithelial cells in flow cytometric analysis, cells were stained with anti-cytokeratin antibody. In addition, the cells were simultaneously stained with anti-p63 and anti-Ki-67 to assess differentiation and proliferation. Positive and negative populations were predefined by compensation setup. Of the cells, 97.02% ± 0.2194% were cytokeratin and p63 positive (Figure 5A), indicating that the isolated cells are undifferentiated epithelial cells. In addition, 86.45% ± 2.532% of cells were Ki-67 positive (Figure 5B), which reflects good proliferation of the cells (Figure 4).

**Figure 5.**
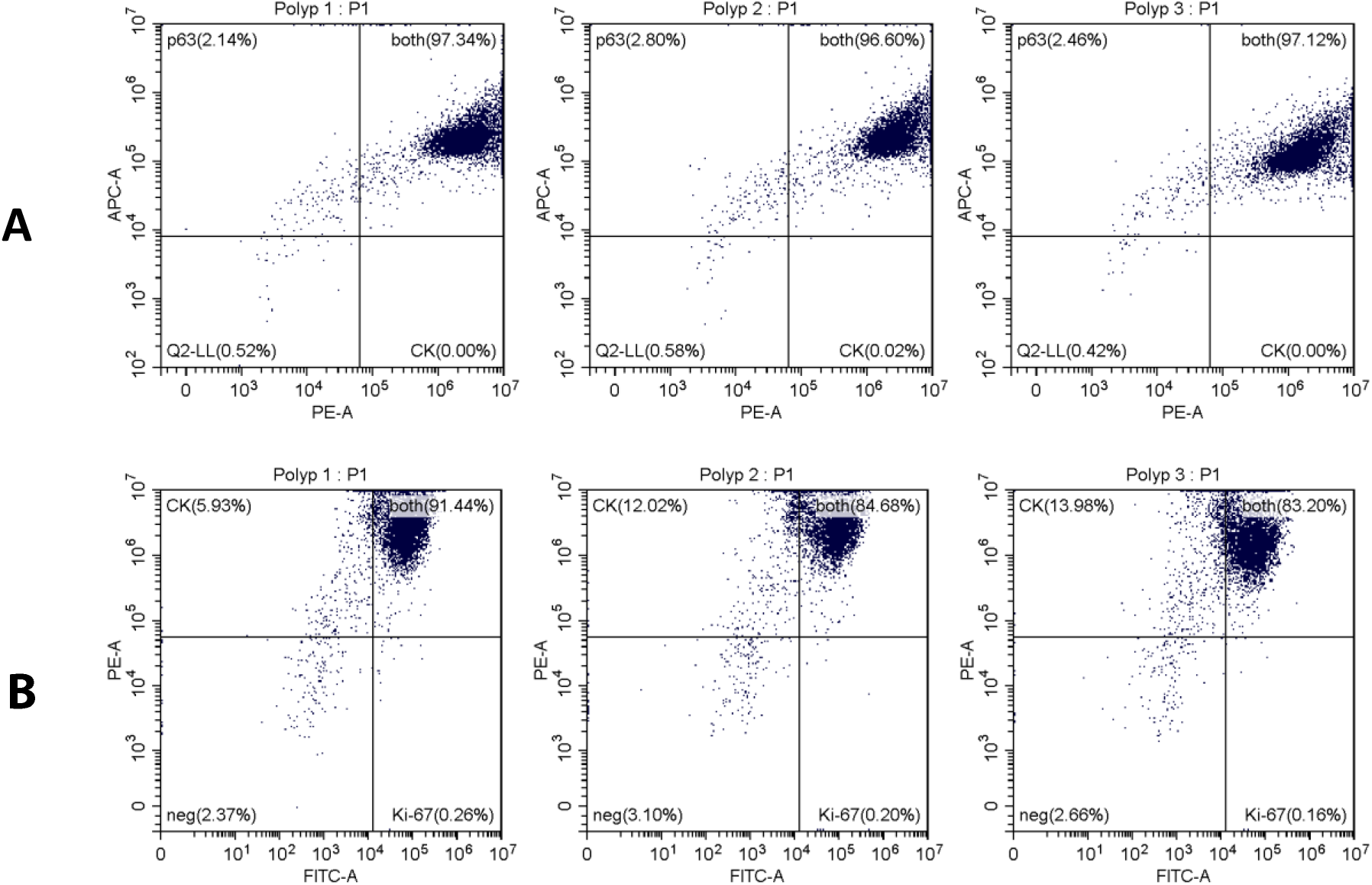
Flow cytometry analysis of cytokeratin, p63, and Ki-67 expression in isolated cells from 3 tissue samples. A: The relative fluorescence intensity due to CK (PE) and p63 (APC) is plotted on the log scale x- and y-axis, respectively. Of the cells, 97.02% ± 0.2194% are cytokeratin and p63 positive, indicating that the isolated cells are undifferentiated epithelial cells. B: The relative fluorescence intensity due to Ki-67 (FITC) and CK (PE) is plotted on the log scale x- and y-axis, respectively. Of the cells, 86.45% ± 2.532% are Ki-67 positive and are therefore in a proliferative cell cycle, which reflects good proliferation of the cells. CK: cytokeratin.

### Functionality test by determining type 2–relevant proteins, using ELISA

The function of the epithelial cells was assessed by determining the production of a pro-inflammatory cytokine and protein produced by the epithelium in type 2 disease, representatively IL-33 and periostin. Cells were able to produce IL-33 (average ± SEM in pg/ml: 21.39 ± 0.7679 in 24 h, 20.18 ± 1.044 in 48 h, and 21.18 ± 1.491 in 72 h; n = 3) and periostin (average ± SEM in pg/ml: 147.5 ± 15.92 in 24 h, 186.5 ± 41.11 in 48 h, and 250.3 ± 18.60 in 72 h, n = 3) (Figure 6), demonstrating sufficient functioning of the epithelial cells for further study.

**Figure 6.**
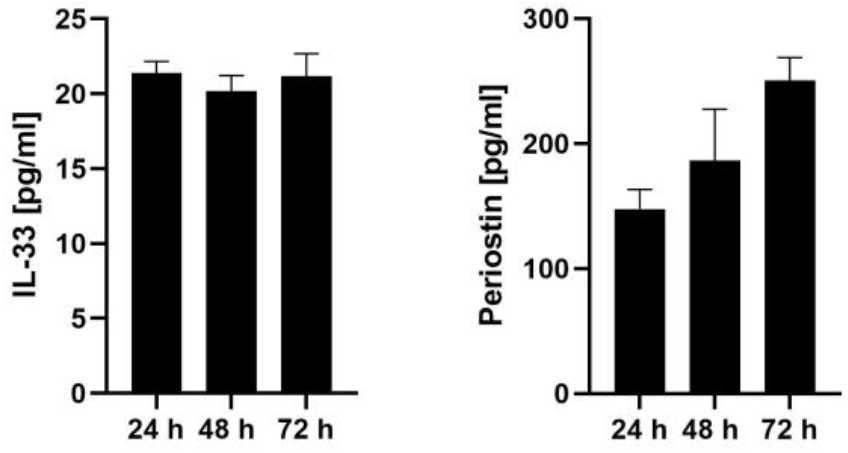
Epithelial cell function was assessed by the production of typical cytokines and proteins of type 2 inflammation, IL-33 and periostin, representatively. Cells were able to produce IL-33 and periostin, demonstrating sufficient functioning of the epithelial cells for further study. IL-33 concentration in pg/ml (n = 3, average ± SEM): 21.39 ± 0.7679 in 24 h, 20.18 ± 1.044 in 48 h, and 21.18 ± 1.491 in 72 h. Periostin concentration in pg/ml (n = 3, average ± SEM): 147.5 ± 15.92 in 24 h, 186.5 ± 41.11 in 48 h, and 250.3 ± 18.60 in 72 h.

## DISCUSSION

So far, the isolation of nasal epithelial cells was performed either with brushing of the mucosa or dissociation of nasal tissue with protease [11,12]. Some researchers even combine the methods [13]. The brushing technique is easy to perform. Particularly, the donation is possible for volunteers who are not undergoing a surgical procedure; however, the ratio of success with this method is approximately 63%–90% [11,14]. The dissociation technique is performed with tissue materials from the surgery. With the dissociation technique, some difficulties are met. An explicit de-epithelization of mucosa is not possible, which differs from, for example, the de-epithelialization of skin tissue, which is possible after incubation with Dispase [12,15]. Therefore, in the mucosa, not only the epithelium is incubated with protease but also the submucosal tissue, which could deteriorate the purity of the cells. A further disadvantage is the possible alteration of the cell physiology by the protease treatment. A study has shown the different DNA methylation profile of bronchial epithelial cells isolated by dissociation with pronase compared to those obtained by the brushing technique [16]. Another disadvantage is that the material cost due to the required proteinase for dissociation, for example, Dispase [12], is significantly higher than with the outgrowth technique.

The outgrowth technique is known as a method for isolating adherent cells, such as fibroblasts [17]. Our outgrowth method showed a good success rate in isolating epithelial cells from nasal polyp tissue (Figure 3). The isolated epithelial cells proliferated well up to the 3rd passage (Figure 4). The purity of the cell population was high (Figure 5A). Almost all of the cells were undifferentiated and active in proliferation (Figure 5A and Figure 5B). The cells were capable of producing the typical inflammatory cytokines and proteins of type 2 inflammation, representatively IL-33 and periostin (Figure 6), which as a pro-inflammatory cytokine trigger a cascade of inflammatory responses [5] and as a marker of the remodeling is able to indicate the intensity of inflammation [18].

In summary, we established a simple, low-cost, and well-performing method for isolating epithelial cells from nasal polyps by using the outgrowth technique.

